# PyBSASeq: a novel, simple, and effective algorithm for BSA-Seq data analysis

**DOI:** 10.1101/654137

**Authors:** Jianbo Zhang, Dilip R. Panthee

## Abstract

Bulked segregant analysis (BSA), coupled with next generation sequencing (NGS), allows the rapid identification of both qualitative and quantitative trait loci (QTL), and this technique is referred to as BSA-Seq here. The current SNP index method and G-statistic method for BSA-Seq data analysis require relatively high sequencing coverage to detect major single nucleotide polymorphism (SNP)-trait associations, which leads to high sequencing cost. Here we developed a simple and effective algorithm for BSA-Seq data analysis and implemented it in Python, the program was named PyBSASeq. Using PyBSASeq, the likely trait-associated SNPs (ltaSNPs) were identified via Fisher’s exact test and then the ratio of the ltaSNPs to total SNPs in a chromosomal interval was used to identify the genomic regions that condition the trait of interest. The results obtained this way are similar to those generated by the current methods, but with more than five times higher sensitivity, which can reduce the sequencing cost by ~80% and makes BSA-Seq more applicable for the species with a large genome.

**Significance Statement:** BSA-Seq can be utilized to rapidly identify DNA polymorphismtrait associations, and PyBSASeq allows the detection of such associations at much lower sequencing coverage than the current methods, leading to lower sequencing cost and making BSA-Seq more accessible to the research community and more applicable to the species with a large genome.

---

Bulked segregant analysis (BSA) has been widely utilized in the rapid identification of trait-associated genetic markers for a few decades (1, 2). The essential part of a BSA study is to construct two bulks of individuals that have contrasting phenotypes (e.g., tallest plants vs. shortest plants or resistant plants vs. susceptible plants) from segregating populations. If a gene does not contribute to the trait phenotype, its alleles would be randomly segregated in both bulks; whereas if a gene is responsible for the trait phenotype, its alleles would be enriched in either bulk, e.g. one bulk has more allele *A* while the other bulk has more allele *a*. BSA was primarily used to develop genetic markers for detecting gene-trait association at its early stage (1, 2). The application of next generation sequencing (NGS) technology to BSA has eliminated the time-consuming and labor-intensive marker development and genetic mapping steps and has dramatically sped up the detection of gene-trait associations (3–20). This technique was termed either QTL-seq or BSA-Seq in different publications (5, 6, 21), we adapted the latter here because it can be applied to study both qualitative and quantitative traits.

The widely used approach in analyzing BSA-Seq data is the SNP index method (5). For each SNP, the base that is the same as in the reference genome is termed reference base (REF), and the other base is termed alternative base (ALT); the SNP index of a SNP is calculated by dividing its ALT read with the total read (REF read + ALT read) in a bulk. The greater the Δ(SNP index) (the difference of the SNP indices between bulks), the more likely the SNP contributes to the trait of interest or is linked to a gene that controls the trait. The second approach is the G-statistic method (21). For each SNP, a G-statistic value is calculated via G-test using the REF read and ALT read values in each bulk. The SNP with a high G-statistic value would be more likely related to the trait. Both methods identify SNP-trait associations via quantifying the REF/ALT enrichment of a single SNP, and some of the major QTLs can be detected only with high sequencing coverage (3, 5, 22), which leads to high sequencing cost and limits the application of BSA-Seq to the species with a large genome.

In BSA studies, bulking enriches the trait-associated alleles in either bulk. The more a gene contributes to the phenotype, the more its alleles are enriched, and so are the SNPs within the gene (one bulk contains more REF read while the other bulk contains more ALT read). The SNPs flanking this gene should be enriched as well due to linkage disequilibrium, the closer the SNP to the gene, the more enrichment is achieved. Such SNPs are termed trait-associated SNPs (taSNPs). We developed a novel, simple, and effective algorithm for analysis of the BSA-Seq data via quantifying the enrichment of trait-associated SNPs in a chromosomal interval. A Python script, PyBSASeq, was written based on this algorithm. The sequence data of Yang *et al.* (3) was used to test our algorithm, and the PyBSASeq method detected more QTLs than the current methods (3, 22) even with lower sequencing coverage.

## Materials and Methods

The sequencing data used in this study were generated by Yang *et al.* (3). Using the G-statistic method, Yang *et al.* identified six major cold tolerance QTLs in rice and five of them were consistent with the then available QTL database or previous publications. The *Oryza sativa* subsp. *japonica* rice cultivar Nipponbare was used as one of the parents in generating the F_3_ population of the BSA-Seq experiment and its genome sequence was used as the reference sequence for SNP calling in our study. The Python implementation of the PyBSASeq algorithm is available on the website https://github.com/dblhlx/PyBSASeq, and its detailed usage can be found on the website as well. The Python implementation of the SNP index method and the G-statistic method can be accessed on https://github.com/dblhlx/.

### SNP calling

The raw sequences (SRR834927 and SRR834931) for BSA-Seq analysis were downloaded from NCBI using fasterq-dump (https://github.com/ncbi/sra-tools) and the sequences were trimmed and quality control was performed using fastp at the default setting (23). The trimmed sequences were aligned to the ‘Nipponbare’ reference genome sequence (Release 41, downloaded from https://plants.ensembl.org/Oryza_sativa/Info/Index) using BWA (24–26). SNP calling was carried out following the best practice of Genome Analysis Toolkit (GATK) (27) and Genome Analysis Toolkit 4 (GATK4) tool documentation on the GATK website https://software.broadinstitute.org/gatk/documentation/tooldocs/current/. The GATK4-generated. vcf file usually contains the information for two bulks; we termed them the first bulk (fb) and the second bulk (sb), respectively. Using the GATK4 tool, the relevant columns (CHROM, POS, REF, ALT, fb.AD, fb.GQ, sb.AD, sb.GQ) of this. vcf file were extracted to create the input file in. tsv (tab separated value) format for the Python script; Table 1 shows the first five rows of this. tsv file.

**Table 1.**
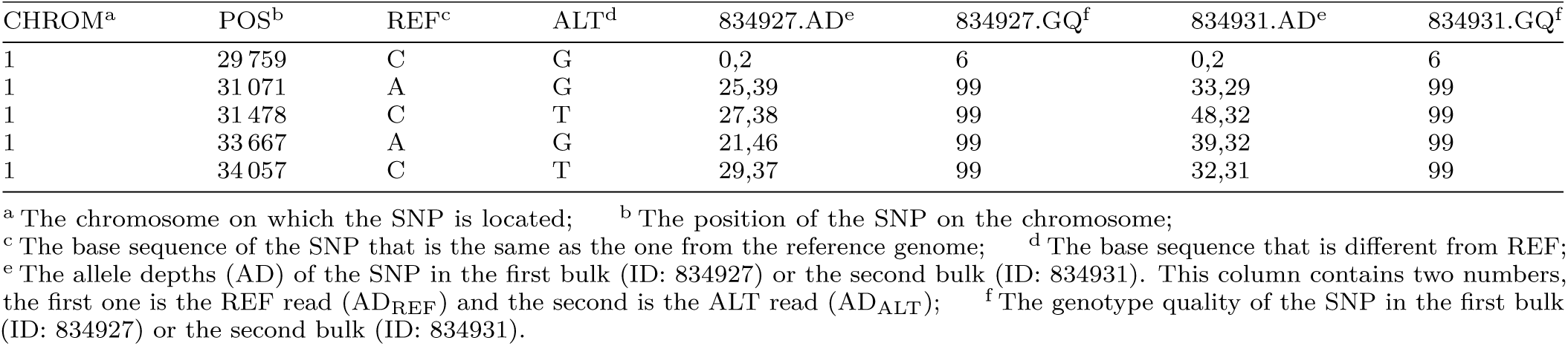
The first five rows of the GATK4 output file

| CHROM<sup>a</sup> | POS<sup>b</sup> | REF<sup>c</sup> | ALT<sup>d</sup> | 834927.AD<sup>e</sup> | 834927.GQ<sup>f</sup> | 834931.AD<sup>e</sup> | 834931.GQ<sup>f</sup> |
| --- | --- | --- | --- | --- | --- | --- | --- |
| 1 | 29 759 | C | G | 0,2 | 6 | 0,2 | 6 |
| 1 | 31 071 | A | G | 25,39 | 99 | 33,29 | 99 |
| 1 | 31 478 | C | T | 27,38 | 99 | 48,32 | 99 |
| 1 | 33 667 | A | G | 21,46 | 99 | 39,32 | 99 |
| 1 | 34 057 | C | T | 29,37 | 99 | 32,31 | 99 |

<sup>a</sup> The chromosome on which the SNP is located; <sup>b</sup> The position of the SNP on the chromosome; <sup>c</sup> The base sequence of the SNP that is the same as the one from the reference genome; <sup>d</sup> The base sequence that is different from REF; <sup>e</sup> The allele depths (AD) of the SNP in the first bulk (ID: 834927) or the second bulk (ID: 834931). This column contains two numbers, the first one is the REF read (AD<sub>REF</sub>) and the second is the ALT read (AD<sub>ALT</sub>); <sup>f</sup> The genotype quality of the SNP in the first bulk (ID: 834927) or the second bulk (ID: 834931).

### SNP filtering

The GATK4-identified SNPs were filtered using the following parameters in order: 1) the unmapped SNPs or SNPs mapped to the mitochondrial or chloroplast genome; 2) the SNPs with a ‘NA’ value in any of the above columns; 3) the SNPs with more than one ALT bases; 4) the SNPs with GQ score less than 20.

### Sliding windows

The sliding window algorithm was utilized to aid the visualization (plotting) in BSA-Seq data analysis. The window size was 2 Mb and the incremental step was 10 000 bp. Empty windows would be encountered if the amount of SNPs is too low or the SNP distribution is severely skewed. If a sliding window has zero SNP, its ltaSNP/totalSNP ratio will be replaced with the value of the previous sliding window. If the first sliding window of a chromosome is empty, the string ‘empty’ will be assigned to this sliding window as a placeholder that will be replaced with a non-empty value of the nearest window later.

### Statistical methods

The number of REF/ALT reads of a SNP is defined as allele depth (AD) in GATK4. Here they are represented as AD_REF_ and AD_ALT_, respectively, and a ‘1’ or ‘2’ is added to its subscript when appropriate to indicate which bulk it belongs to; the same can be applied to the sequencing depth as well. In some rare occasions, the GATK4-generated depth per sample (DP) of a SNP can be greater or less than the sum of the ADs in a bulk, here DP was defined as below for all the SNPs:

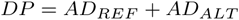

#### Fisher’s exact test

Python module scipy.stats.fisher_exact(ctbl) was used for Fisher’s exact test; *ctbl* is a 2×2 contingency table comprising all the AD values of a SNP in both bulks and is represented as a numpy array ([[AD_REF1_, AD_ALT1_], [AD_REF2_, AD_ALT2_]]). This module returns a pair of numbers, and the second number is the p-value of the Fisher’s exact test.

#### Calculation of G-statistic

The following formula was used for this purpose, where *O* is the observed AD, *E* is the expected AD under the null hypothesis, and ln denotes the natural logarithm.

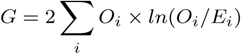

#### Simulation of AD_REF_/AD_ALT_ for threshold calculation

The python module numpy.random.binomial(DP, alleleFreq) was used to calculate the simulated AD_REF_ (smAD_REF_) and simulated AD_ALT_ (smAD_ALT_) of a SNP in a bulk; *alleleFreq* is the frequency of the ALT base in the bulk under the null hypothesis that the SNP is not associated with the trait, and its value was obtained via simulation (see the *smAlleleFreq* function of the Python script for details). The module returns the smAD_ALT_, and the smAD_REF_ can be calculated as below:

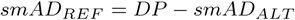

#### Calculation of the Δ(SNP index) and G-statistic thresholds

For each SNP in the SNP dataset, smAD_REF1_/smAD_ALT1_ of bulk 1 and smAD_REF2_/smAD_ALT2_ of bulk 2 were obtained as described above. Using these AD values, the. Δ(SNP index) was calculated with the equation below and the G-statistic was calculated as previously stated. This process was repeated 10 000 times, the 99% confidence interval of the 10 000. Δ(SNP index) values was used as the significant threshold for the SNP index method, and the 99.5^th^ percentile of the 10 000 G-statistic values was used as the significant threshold for the G-statistic method.

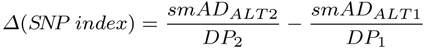

## Results

### Identify SNPs likely associated with the trait of interest

In BSA-Seq studies, each bulk contains many individuals that could be either homozygous (REF or ALT) or heterozygous in any SNP locus, and the bulk is collectively sequenced. Hence most SNPs identified via the SNP calling pipeline contains both the reference base (REF) and the alternative base (ALT) in each bulk. Due to phenotypic selection via bulking, the REF/ALT base of a trait-associated SNP would be enriched in either bulk, and the ALT (or REF) read proportions should be significantly different between the bulks. Fisher’s exact test was performed to identify such SNPs using the AD_REF_ and AD_ALT_ of each SNP in both bulks. A small p-value of the Fisher’s exact test suggests that the ALT proportion difference of a SNP between bulks is more likely caused by bulking and a SNP with its p-value less than 0.01 was considered more likely associated with the trait and was termed ltaSNP here. 240 351 ltaSNPs were identified among total 1 303 084 filtered SNPs (see materials and methods section for the filter criteria), and the chromosomal distribution of SNPs was summarized in Table 2. The chromosomes 8, 1, 2, 10, and 5 contained the most ltaSNPs and had the highest ltaSNP/totalSNP ratios, correlating perfectly with the chromosomes carrying the verified QTLs (3, 22).

**Table 2.** Chromosomal distribution of SNPs

|  | ltaSNP | totalSNP | ltaSNP/totalSNP |
| --- | --- | --- | --- |
| 1 | 52 093 | 160 780 | 0.324 |
| 2 | 48 912 | 125 059 | 0.391 |
| 3 | 3502 | 45 927 | 0.076 |
| 4 | 3743 | 62 317 | 0.060 |
| 5 | 15 482 | 102 474 | 0.151 |
| 6 | 7653 | 159 857 | 0.048 |
| 7 | 12 679 | 128 658 | 0.099 |
| 8 | 54 372 | 132 646 | 0.410 |
| 9 | 1709 | 57 971 | 0.029 |
| 10 | 28 711 | 98 646 | 0.291 |
| 11 | 5235 | 180 319 | 0.029 |
| 12 | 6260 | 48 430 | 0.129 |
| Genome-wide | 240 351 | 1 303 084 | 0.184 |

### Enrichment of ltaSNPs

The ltaSNPs should cluster around the genes controlling the trait phenotype on the chromosomes due to linkage disequilibrium. Using the sliding window technique, the number of ltaSNPs was plotted across all the chromosomes to test if this was the case. We found the ltaSNP plot approximately matched with the major peaks in plots produced by the SNP index method and the G-statistic method (3, 22) (Figure 1A). However, counting the absolute number of ltaSNPs is not an ideal way to measure the ltaSNP enrichment because SNPs were distributed unevenly across and between chromosomes (Figure 1A); if a gene that conditions the trait is located in a region with fewer SNPs, it would be missed using this approach. Thus, we used the ratio of ltaSNPs to total SNPs in a chromosomal region to measure the ltaSNP enrichment. The ltaSNP/totalSNP ratios were plotted for all the chromosomes (Figure 1B), and the plot pattern matched very well with that produced by the G-statistic method (3, 22). The most obvious difference between Figure 1A and figure 1B was the first peak on chromosome 2 and the peaks on chromosomes 3, 6 and 9; these regions contained fewer SNPs, but the ltaSNPs enrichment was relatively high.

**Figure 1.**
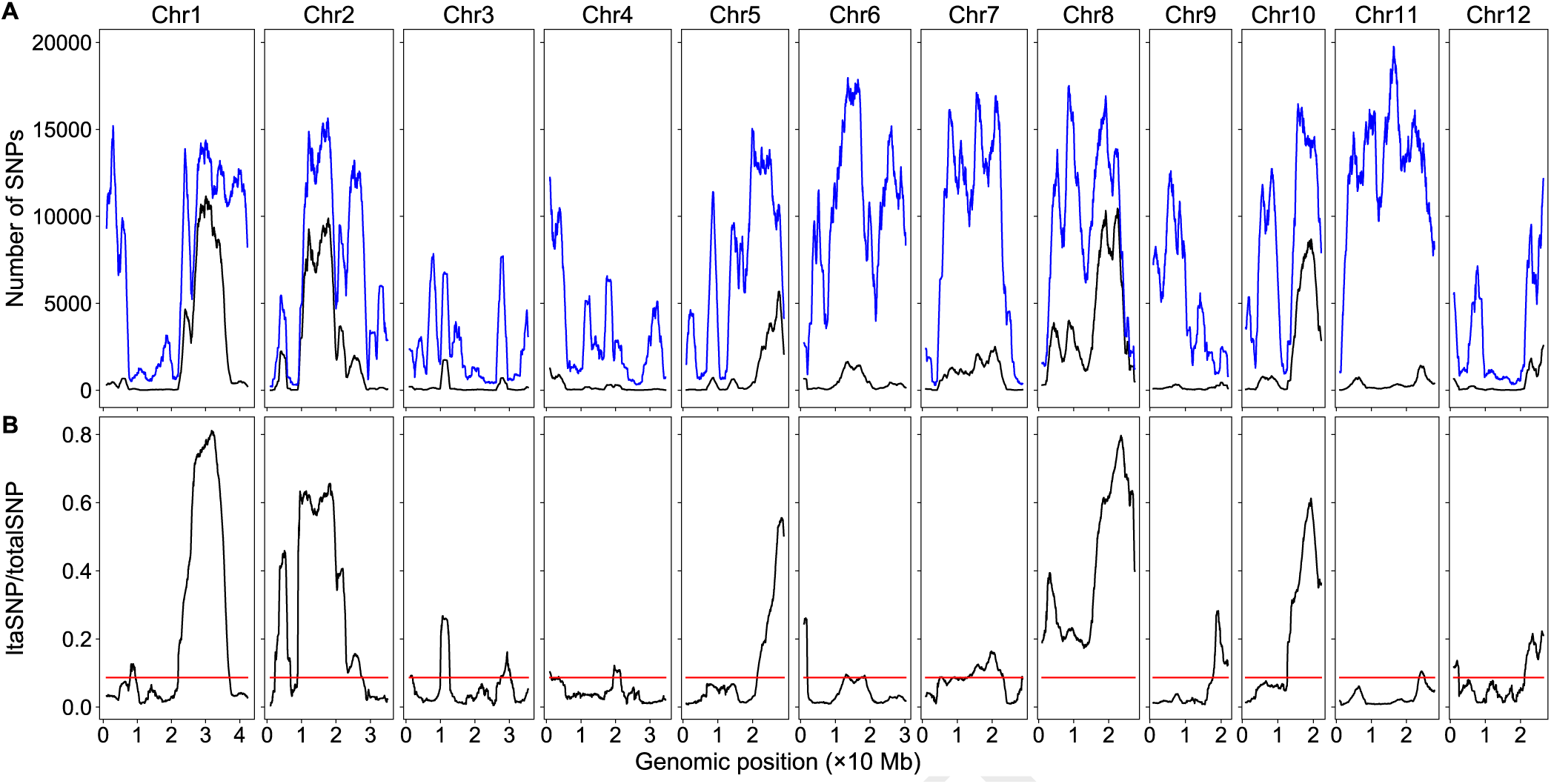
Genomic distributions of SNPs and ltaSNPs/totalSNP ratios. The red horizontal lines are the thresholds. **(A)** The ltaSNPs (black) and total SNPs (blue). **(B)** The ratio of ltaSNPs and total SNPs.

Under the null hypothesis that the SNPs were not associated with the trait, resampling was utilized to obtain the threshold to determine which peak in Figure 1B was statistically significant. For each SNP in the dataset, its simulated AD_REF_ and AD_ALT_ were calculated as detailed in the materials and methods section and then the simulated AD_REF_ and AD_ALT_ from both bulks were used to perform Fisher’s exact test. A SNP with its p-value less than 0.10 was considered a ltaSNP (A high cut-off p-value results in a high threshold). The amount of SNPs that are the same as the average number of SNPs per sliding window were randomly selected from the SNP dataset and the simulated ltaSNP/totalSNP ratio (total SNPs was the sample size) in the sample was recorded. This process was repeated 10 000 times, and the 99.5^th^ percentile of these 10 000 values was used as the significant threshold for the detection of peak-trait associations. The threshold obtained this way was 0.087. In addition to the six major QTLs (two of them on chromosome 2) verified in the work of Yang *et al.* (3), one or more new peaks on all chromosomes except chromosomes 5 and 10 were also above the threshold (Figure 1B).

### Sequencing coverage affected the detection of SNP-trait association

Using the Lander/Waterman equation (28), the sequencing coverage of SRR834927 and SRR834931 was estimated to be 84× and 103×, respectively. It would be very costly to achieve such high sequencing coverage for the organisms with a large genome. Thus, we wanted to know how decreasing sequencing coverage would affect the detection of SNP-trait associations. To achieve lower sequencing coverage, we sampled 40%, 30%, and 20% of the raw sequence reads using the seqtk program (https://github.com/lh3/seqtk) with random seeds 123, 160, and 100, respectively. The ltaSNPs were identified from these sequence subsets and the ratios of ltaSNP/totalSNP were plotted along all the chromosomes as above. The results revealed that the plotting patterns were very similar at different sequencing coverage levels (Figure 2); with decreasing sequencing coverage, the total SNPs decreased slightly, while the number of ltaSNP and the ltaSNP/totalSNP ratio decreased substantially (Table S1). Because the threshold did not change as much, more and more minor SNP-trait associations were missed with decreasing sequencing coverage. However, with 40%, 30%, or even 20% of the original sequencing coverage, more QTLs were detected than the current methods with the original sequencing coverage (3, 22).

**Figure 2.**
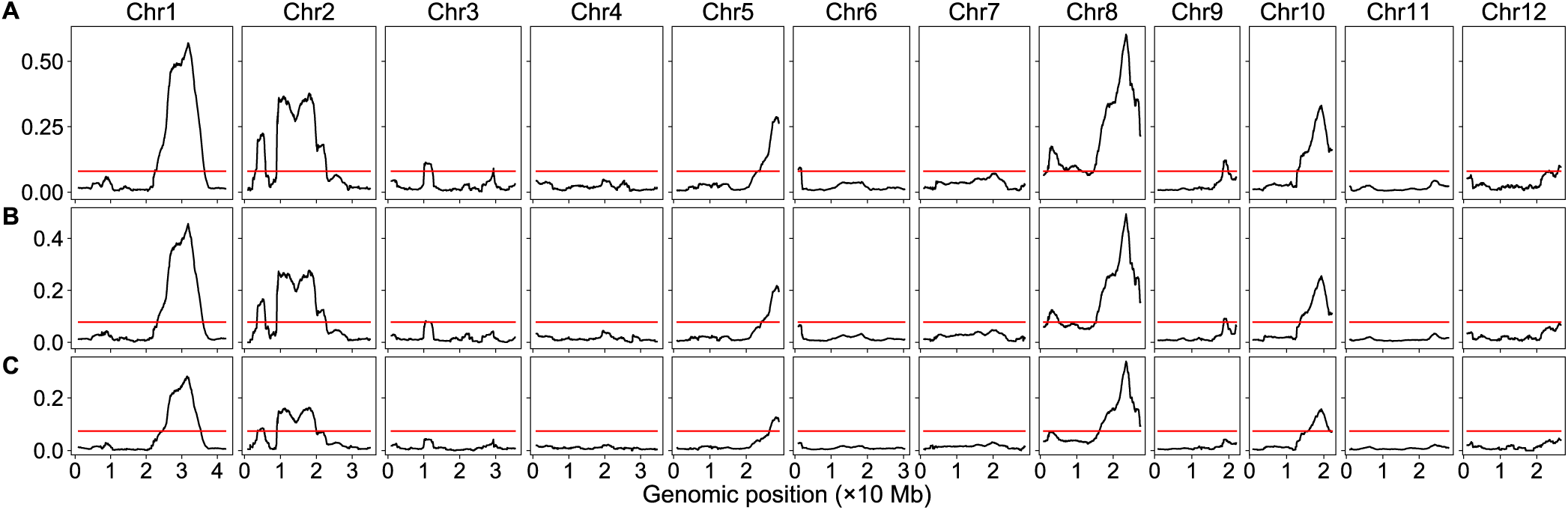
Genomic distribution of ltaSNP/totalSNP ratios at different sequencing coverage levels. The red horizontal lines are the thresholds. **(A)** 40% of the original sequence reads. **(B)** 30% of the original sequence reads. **(C)** 20% of the original sequence reads.

### Sensitivity comparison

The above data indicated that the PyBSASeq approach had higher detection power. However, different methods were used to generate the SNP datasets (3, 22), which might lead to different detection sensitivities. To rule out this possibility, we implemented the SNP index method and the G-statistic method in Python and tested all the three methods with the same SNP dataset. First, we tested if the results of Yang *et al.* and Mansfeld and Grumet can be replicated using our scripts. As in the studies mentioned above, the SNP dataset was filtered with the following criteria: fb.GQ ≥ 99, sb.GQ ≥ 99, fb.DP ≥ 40, sb.DP ≥ 40, fb.DP+sb.DP ≥ 100, and fb.DP+sb.DP ≤ 400. Although the SNP datasets were generated in different ways (GATK4 vs. GATK vs. Samtools) and no smoothing besides the sliding window algorithm was applied in our scripts, the results, including the plot patterns, the G-statistic values, and the Δ(SNP index) values and its confidence intervals, were very similar (3, 22), and the positions of the peaks/valleys matched almost perfectly between different approaches (Figure S1). A non-parametric method was used to calculate the threshold in the G-statistic method by Yang *et al.* and Mansfeld and Grumet, and different approaches were used to remove the G-statistic values from the QTL regions. Thus the thresholds were a little different in these studies and so was the QTL detection results (3, 22). In our G-statistic script, we used simulation for threshold calculation (see the materials and methods), and the thresholds obtained this way were consistent across all the chromosomes and was less conservative than the previously reported approaches. Using the high sequencing depth SNP subset, similar results were obtained by both the SNP index method and the G-statistic method: the six major QTLs and a minor QTL on chromosome 2 were detected (Figure S1). However, the PyBSASeq approach had the highest sensitivity using the same filtering criteria, and it can detect more minor QTLs than other methods even if the whole SNP dataset was used (Figures 1B, 2, and S1).

As in PyBSASeq, we also tested how decreasing sequencing coverage would affect the detection of the SNP-trait associations in these two methods. Using the original sequencing reads, the SNP index method had relatively low detection power, the major QTL on chromosome 5 was missed and the peak (valley) representing the major QTL on chromosome 10 was barely beyond the threshold. With decreasing sequencing coverage, the Δ(SNP index) did not change much, but the thresholds increased dramatically, the QTLs on chromosomes 2, 5, and 10 were missed at 40% of the original sequencing coverage and all the QTL were missed at 30% or lower of the original sequencing coverage (Figure 3). For the G-statistic method, with the original sequencing reads, all the 6 major QTLs can be detected. With decreasing sequencing coverage, the G-statistic values decreased substantially, whereas the threshold increased slightly; the QTLs on chromosomes 2, 5, and 10 were missed at 40% of the original sequencing coverage, the peaks representing the QTLs on chromosomes 1 and 8 were barely above the threshold at 30% of the original sequencing coverage, and all the QTLs were missed at 20% of the original sequencing coverage (Figure 4).

**Figure 3.**
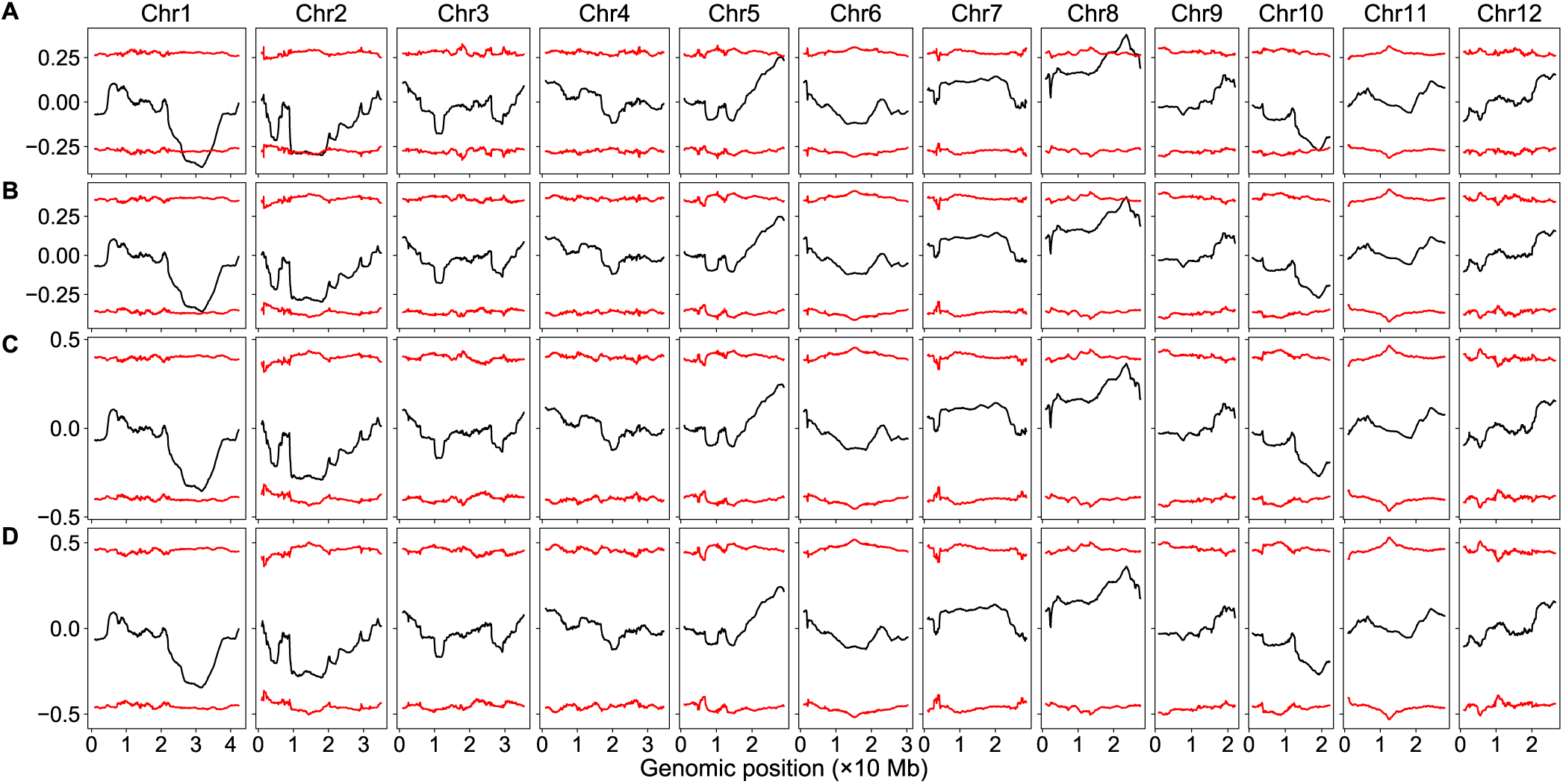
Genomic distribution of. 6. (SNP index) at different sequencing coverage levels. The red curves indicate the 99% confidence intervals. **(A)** The original sequence reads. **(B)** 40% of the original sequence reads. **(C)** 30% of the original sequence reads. **(D)** 20% of the original sequence reads.

**Figure 4.**
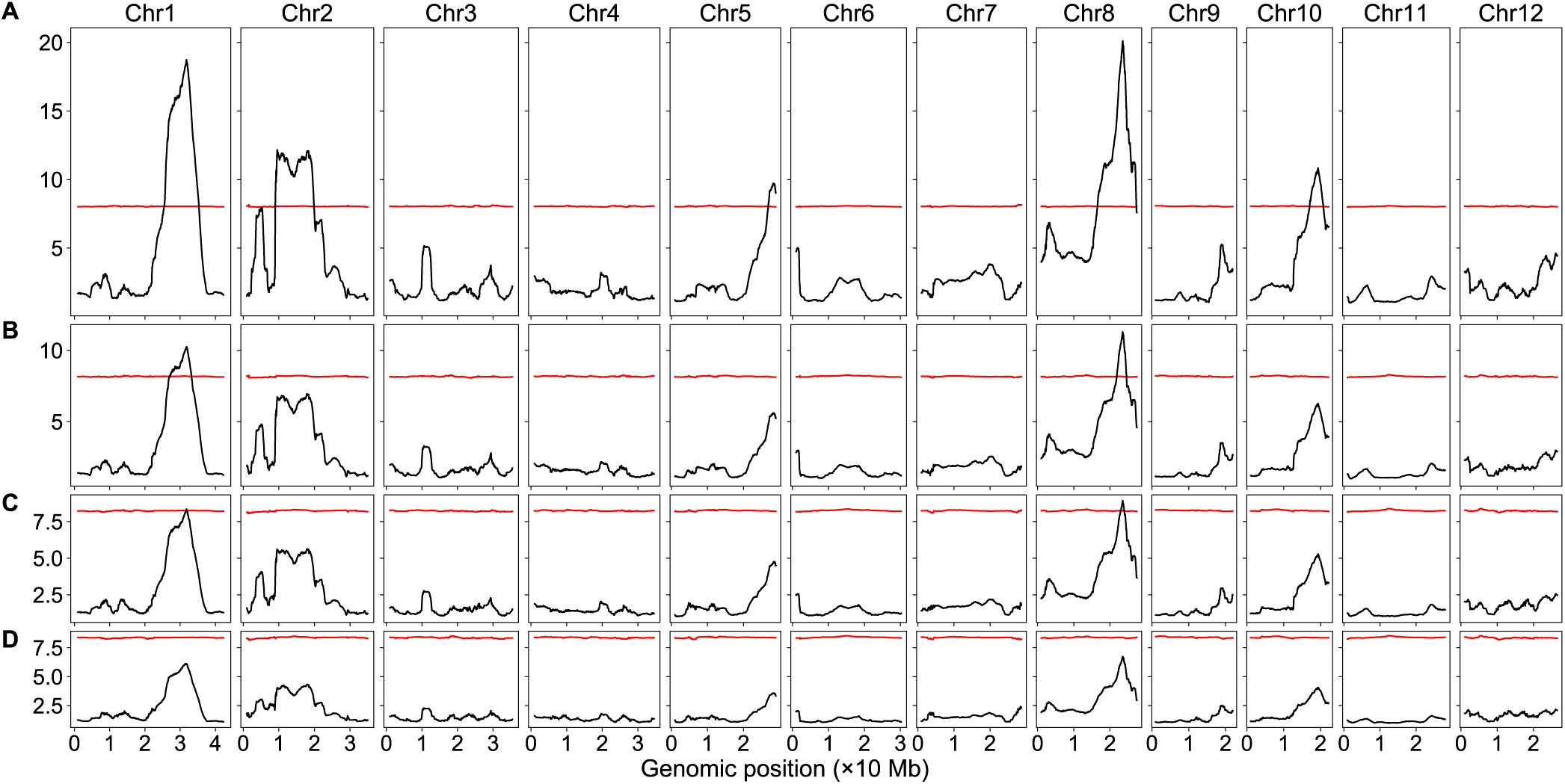
Genomic distribution of G-statistic at different sequencing coverage levels. The red curves are the G-statistic thresholds. **(A)** The original sequence reads. **(B)** 40% of the original sequence reads. **(C)** 30% of the original sequence reads. **(D)** 20% of the original sequence reads.

## Discussion

PyBSASeq detected more than 10 minor QTLs along with all of the major QTLs detected via the current methods when run with the entire SNP dataset based on the original sequencing reads (Figures 1B, 3A, and 4A). Plant cold tolerance is a complex quantitative trait controlled by many genes (29, 30). The additional QTLs detected via PyBSASeq may represent the minor QTLs that have small phenotypic effects. Filtering out the SNPs with a low DP value increased the sensitivity of the current methods (Figures S1, 3, and 4), but doing so increased the sensitivity of PyBSASeq as well (Figures S1C and 1B). Decreasing the sequencing coverage substantially reduced the detection power of all the methods (Figures 2, 3, and 4). At 20% of the original coverage (17× in the first bulk and 21× in the second bulk) all QTLs were missed using the current methods; however, all the verified major QTLs plus two minor QTLs can still be detected via PyBSASeq, manifesting that PyBSASeq is at least five times more sensitive.

Because of its high sensitivity, the intervals of the QTLs (chromosomal regions above the threshold) are quite wide (Figure 1). An extreme case is chromosome 8 where all of its ltaSNP/totalSNP ratios are greater than the threshold, which does not imply that all the SNPs on chromosome 8 are involved in conditioning the cold tolerance trait. The SNPs in the causal locus are enriched because of phenotypic selection via bulking while the SNPs flanking the causal locus are enriched because of linkage disequilibrium. Any recombination event between the SNPs that affects the trait of interest and the SNPs flanking the causal gene would reduce the enrichment of the flanking ltaSNPs, thus SNPs in the causal locus should have the highest enrichment and should be located in the peak region. Therefore, there are only two QTLs on chromosome 8: a minor one on the proximal arm while a major one on the distal arm of the chromosome. All three methods use the sliding window algorithm to detect the SNP-trait associations and should have the same level of resolution if the sliding window settings (window size and incremental step) are the same.

Both the SNP index method and the G-statistic method identify SNP-trait associations by measuring REF/ALT enrichment of a single SNP; whereas the PyBSASeq method identifies SNP-trait associations by measuring ltaSNP enrichment in a chromosomal region. The average number of SNPs was 6984 in a sliding window, much higher than the average sequencing coverage in either bulk (84× in the first bulk and 103× in the second bulk), which could be why PyBSASeq has much higher statistical power. GATK is widely used for SNP and small InDel calling and the new version of GATK4 is also capable of copy number and structural variant calling. PyBSASeq is designed to analyze the GATK-generated variant calling data, though it has only been tested for analysis of the SNP and small InDel calling data, it should be able to handle the GATK4-generated copy number variant and structural variant data as well.

## Conclusions

The high sensitivity of PyBSASeq allows the detection of SNP-trait associations at reduced sequencing coverage, leading to reduced sequencing cost. Thus, BSA-Seq can be more practically applied to species with a large genome.

## Author contributions

JZ and DRP conceived the study. JZ developed the algorithm, wrote the Python code, analyzed the data, and wrote the manuscript. DRP edited the manuscript and supervised the project.

The authors declare no conflict of interest.

## Acknowledgments

JZ is supported by the National Science Foundation grant [IOS-1546625 to DRP]. We are grateful to Yang *et al.* for generating the sequencing data and making it available to the public. We thank Dr. Thomas Ranney and Nathan Lynch for valuable comments. The manuscript was prepared using a modified version of the PNAS L^A^T_E_X template.

## Supplementary Information

**Table S1.** Chromosomal distribution of SNPs at different sequencing coverage levels

| Chromosome | 40% of the original coverage |  |  | 30% of the original coverage |  |  | 20% of the original coverage |  |  |
| --- | --- | --- | --- | --- | --- | --- | --- | --- | --- |
|  | ltaSNP | totalSNP | ltaSNP/totalSNP | ltaSNP | totalSNP | ltaSNP/totalSNP | ltaSNP | totalSNP | ltaSNP/totalSNP |
| 1 | 28 501 | 150 662 | 0.189 | 20 464 | 144 815 | 0.141 | 11 122 | 133 953 | 0.083 |
| 2 | 23 760 | 116 704 | 0.204 | 16 531 | 111 944 | 0.148 | 8 727 | 103 470 | 0.084 |
| 3 | 1492 | 43 650 | 0.034 | 1030 | 42 307 | 0.024 | 592 | 39 406 | 0.015 |
| 4 | 1628 | 59 037 | 0.028 | 1127 | 56 702 | 0.020 | 674 | 52 609 | 0.013 |
| 5 | 6865 | 96 583 | 0.071 | 4892 | 93 201 | 0.052 | 2776 | 86 864 | 0.032 |
| 6 | 3302 | 150 246 | 0.022 | 2404 | 145 069 | 0.017 | 1361 | 134 562 | 0.010 |
| 7 | 5122 | 120 045 | 0.043 | 3549 | 115 430 | 0.031 | 1925 | 106 121 | 0.018 |
| 8 | 26 933 | 123 411 | 0.218 | 19 378 | 118 699 | 0.163 | 10 652 | 108 581 | 0.098 |
| 9 | 825 | 55 088 | 0.015 | 680 | 53 291 | 0.013 | 424 | 49 768 | 0.009 |
| 10 | 13 039 | 92 815 | 0.140 | 9477 | 89 639 | 0.106 | 5526 | 82 996 | 0.067 |
| 11 | 2622 | 171 170 | 0.015 | 1963 | 165 570 | 0.012 | 1313 | 154 588 | 0.008 |
| 12 | 2589 | 46 061 | 0.056 | 1775 | 44 596 | 0.040 | 995 | 41 051 | 0.024 |
| Genome-wide | 116 678 | 1 225 472 | 0.095 | 83 270 | 1 181 263 | 0.070 | 46 087 | 1 093 969 | 0.042 |

**Figure S1.**
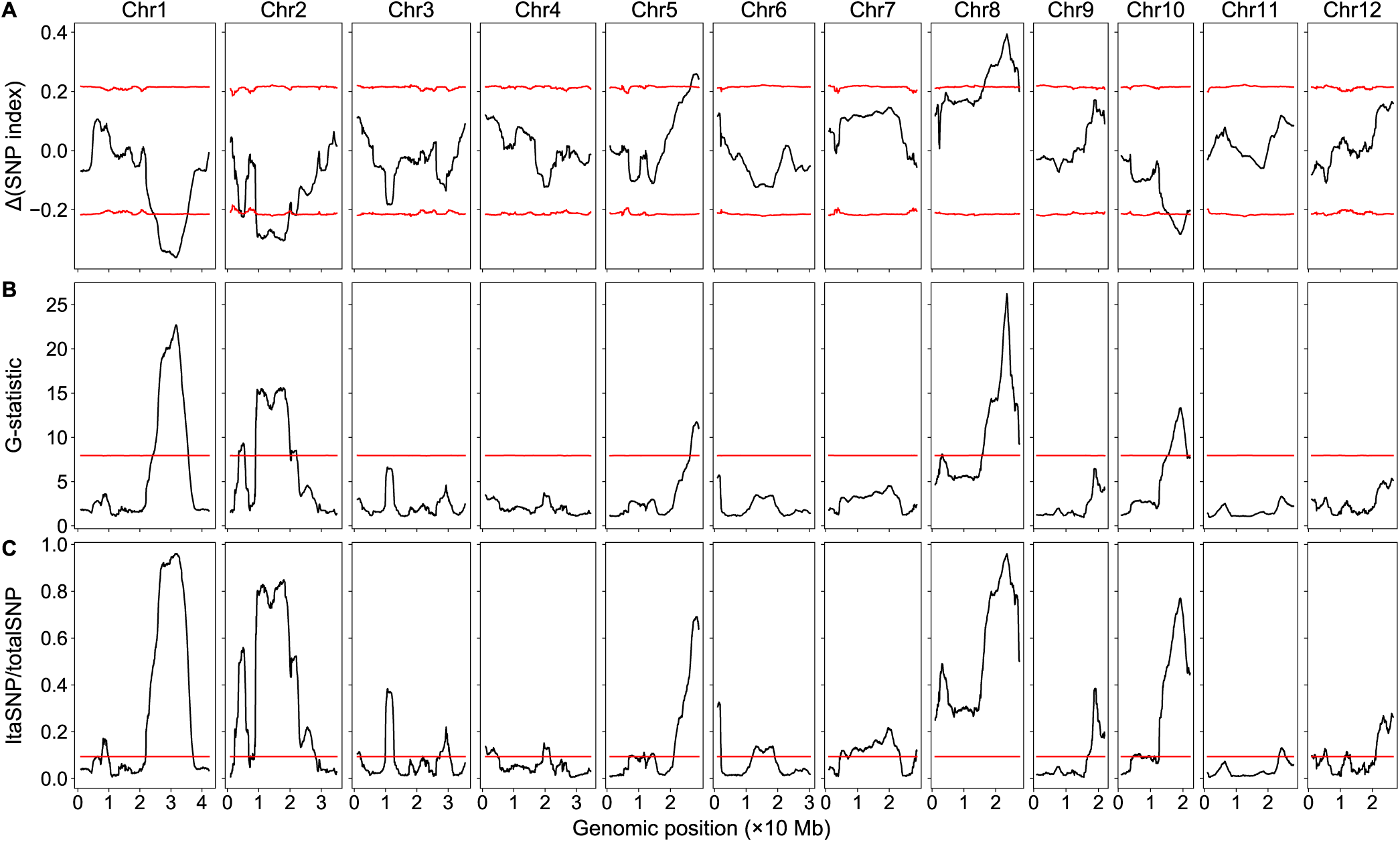
Replication of the SNP index method and the G-statistic method in Python. **(A)** The SNP index method. The red curves indicate the 99% confidence intervals. **(B)** The G-statistic method. The red curves indicate the G-statistic thresholds. **(C)** The PyBSASeq method included here for comparison. The red lines are the thresholds.

